# Microbial species exist and are maintained by ecological cohesiveness coupled to high homologous recombination

**DOI:** 10.1101/2024.05.25.595874

**Authors:** Roth E. Conrad, Catherine E. Brink, Tomeu Viver, Luis M. Rodriguez-R, Borja Aldeguer-Riquelme, Janet K. Hatt, Stephanus N. Venter, Rudolf Amann, Ramon Rossello-Mora, Konstantinos T. Konstantinidis

## Abstract

Recent analyses of metagenomes and genomes have revealed that microbial communities are predominantly composed of persistent, sequence-discrete species and intraspecies units (genomovars). To advance the species concept the underlying genetic or ecological mechanisms that maintain these discrete units need to be elucidated. By analyzing closely related isolate genomes from the same or related samples we show that high ecological cohesiveness coupled to frequent-enough and unbiased (i.e., not selection driven) horizontal gene flow, mediated by homologous recombination, often underlie these diversity patterns. Ecological cohesiveness was inferred based on higher similarity in abundance patterns of genomes of the same vs. different units, while recombination frequency was shown to have two times or more impact on sequence evolution than point mutation. Therefore, our results represent a departure compared to previous models of microbial speciation that invoke either ecology or selection-driven recombination, but not their synergistic effect, as the mechanism of unit cohesion. These results were observed in both *Salinibacter ruber*, an environmental halophilic organism, and *Escherichia coli*, the model gut-associated organism and an opportunistic pathogen, indicating that they may be more broadly applicable to the microbial world. Therefore, our results have strong implications for how to identify and regulate microbial species and genomovars of clinical or environmental importance and answer an important question for microbiology: what a species is.

**SIGNIFICANCE:** A highly pressing issue to resolve toward advancing the species concept for microbes (i.e., “what a species is”) is to elucidate the underlying mechanisms for creating and maintaining species- and intraspecies-level gaps in diversity, or simply “clusters”. In this study, we provide a novel methodology and the appropriate data to elucidate these mechanisms, and thus provide a mechanistic explanation of how the evolution of species- and strain-level clusters takes place. Specifically, our results show that several bacteria may be evolving and speciating much more sexually than previously thought, even under conditions of no strong positive selection for DNA exchange (i.e., neutral conditions). These results have major implications for better understanding and modeling microbial diversity on the planet.

## INTRODUCTION

Whether species exist and, if so, how to recognize them are challenging questions to answer for many microbes including Bacteria and Archaea (the prokaryotes), with obvious practical implications for identifying or regulating organisms of clinical or environmental importance (Gevers 2005, Konstantinidis 2006, Rossello-Mora and Amann 2015, Konstantinidis 2023). Recent large-scale surveys of prokaryotic communities (metagenomes) as well as isolate genomes have revealed that their diversity is predominantly organized in sequence-discrete clusters or units that may be equated to species. Specifically, genomes of the same species commonly show average nucleotide identity (ANI) of shared genes >95% between them and ANI <85% to members of other species (Caro-Quintero and Konstantinidis 2012, Bendall 2016, Olm 2020, Rodriguez 2021). Intermediate identity genotypes, for example, sharing 85–95% ANI, when present, are generally ecologically differentiated and scarcer in abundance, and thus should probably be considered distinct species (Viver 2020, Rodriguez 2021) rather than representing cultivation or other sampling biases (Murray 2021). Sequence-discrete clusters similar to those described above for prokaryotes have recently been recognized for eukaryotic protozoa (Seabolt 2021) and different types of viruses, including bacteriophages and viruses of eukaryotic hosts (Simmonds 2017, Roux 2021). Therefore, it appears that similar species-level diversity patterns may characterize most microbes and viruses.

More recently, our team observed another discontinuity (or gap) in ANI values that may be used to define the units within a species, most notably genomovars and strains (Rodriguez 2023, Viver 2024). Specifically, the analysis of all complete isolate genomes (n=18,123) from 330 diverse bacterial species revealed a clear bimodal distribution in the ANI values within the great majority (>95%) of these species. That is, there is a scarcity of genome pairs showing 99.2-99.8% ANI (midpoint at 99.5% ANI) in contrast to genome pairs showing ANI >99.8% or <99.2% (Rodriguez 2023). We also suggested that the term genomovar could be used to refer to these 99.5%-ANI intraspecies units. We did not observe another pronounced ANI gap within the 99.5% ANI clusters, and thus recommended the use of the term strain only for nearly-identical genomes based on the prevailing expectation that members of the same strain should be phenotypically very similar. Specifically, we proposed to define a strain as a collection of genomes sharing ANI >99.99% based on the high gene-content similarity observed among genomes at this high ANI level, e.g., typically, >99.0% gene content is shared (Viver 2024). These definitions are largely consistent with how several units within species such as sequence types and strains have been recognized previously, but provide units that encompass genomically more homogenous organisms compared to the existing practice and the means to standardize intraspecies definitions across taxa (Rodriguez 2023). Accordingly, we use these definitions below for genomovars and strains.

To better describe and model these diversity patterns, it is imperative to understand what mechanisms underlie the maintenance of sequence-discrete species and genomovars; that is, how members of a sequence-discrete unit cohere together. Several competing hypotheses based on functional differentiation (*ecological species*), recombination frequency (*recombinogenic species*), or variations of these hypotheses, have been advanced to explain the 95% ANI species gap (or the newer 99.5% ANI gap within species) [reviewed in (Fraser 2007, Shapiro and Polz 2015, Konstantinidis 2023)]. Specifically, ecological speciation includes cases in which members of the same species could be functionally differentiated from members of other, related species (or genomovars of the same species) either due to specialization for different growth conditions or different affinities for the same energy substrate. Selection over time for these functions favors the growth of the corresponding members and may result in purging (loss) of diversity (e.g., loss of members not carrying these functions) and thus speciation. Notably, given an estimated mutation rate of ∼4×10^-10^ per nucleotide per generation (Drake 1998) and between 100 to 300 generations per year (Gibbons 1967), it would take two distinct lineages of a gut microbe such as *Escherichia coli* at least 100,000 years since their last common ancestor to accumulate 0.5% difference (i.e., fixed mutations) in their core genes or 99.5% ANI. Therefore, given enough time, it is possible to have ecological purging of diversity even at around the 99.5% ANI level (let alone at 95% ANI) that accounts for the ANI patterns observed. While a few examples of ecological speciation have recently been reported for prokaryotes based -primarily- on differential use of growth substrates (Chase 2021, Strachan 2023), these were not directly associated with the prevailing sequence-discrete species recovered by the metagenomes. That is, these studies have shown that ecological speciation is possible but to what extent it accounts for the sequence-discrete units remains to be evaluated. Further, these studies typically involved laboratory enrichment studies with strong selection pressures (e.g., high substrate concentrations), which may be rather different compared to natural conditions.

Members of a species (or a genomovar) could cohere together via means of unbiased (random) gene exchange which is more frequent within than between species. [Note that this is a fundamentally different mechanism than the sexual reproduction in eukaryotes and the accompanying biological species concept, although the ultimate outcome in terms of species cohesion may be similar. That is, gene exchange does not occur during a meiosis step but via vectors of horizontal gene transfer mediated by recombination. Hence, we opted to not use biological or sexual species here and, instead, refer to them as recombinogenic species]. Indeed, several studies have concluded that the frequency of homologous recombination could be high enough to have a greater effect on sequence evolution than point (or diversifying) mutation, as has been shown to be the case for *Campylobacter* species (Sheppard 2008) and other taxa more recently (Bobay and Ochman 2017). However, these studies have not been able to assess whether recombination is occurring across the genome (that is, it is not biased spatially) to serve as a force of cohesion. In fact, at least in the *Campylobacter* case, recombination was convincingly shown to be biased to a few specific regions (genomic islands) of the genome and functions (e.g., antibiotic resistance and motility) that are apparently under strong positive selection. Thus, recombination is unlikely to lead to species cohesion in such cases since the non-recombining segments of the genome will continue to diverge (Caro-Quintero 2009). In summary, the question of whether or not recombination is frequent and random enough across the genome to maintain the sequence-discrete units remains to be more rigorously tested. By analyzing available closely related genomes here we show that recombination frequency coupled to ecological cohesiveness among members of these units might account for the species and sub-species clusters observed previously.

## RESULTS AND DISCUSSION

To obtain new insights into the role of recombination as a force of species/unit cohesion, we focused initially on a well-sampled bacterial species, *Salinibacter ruber (Sal. ruber)*, which thrives in natural or engineered hypersaline environments, and subsequently evaluated the applicability of the resulting findings with a recently reported collection of *Escherichia coli* isolate genomes originating within a ∼100 km radius from livestock farms in the United Kingdom (Shaw 2021). Engineered solar salterns are operated in repeated cycles of feeding with natural saltwater, increasing salt concentration due to water evaporation caused by natural sunlight, and finally, salt precipitation for human consumption. Previous studies have shown that salterns in different parts of the world harbour recurrent microbial communities each year (Casamayor 2002, Gomariz 2015). These communities show low class/family diversity, generally consisting of two major lineages i.e., the archaeal *Halobacteria* class and the bacterial family of *Salinibacteraceae*, class *Rhodothermia* (Anton 2000, Gomariz 2015, Mora-Ruiz 2018), but with relatively high species richness within each class (Viver 2018, Viver 2019). Notably, *Sal. ruber* makes up at least 1- 2% of the total microbial community in salterns in any sample characterized to date, and typically between 5-25% of the total; that is, it represents a highly abundant member of the saltern communities and it is easy to isolate in pure culture (Viver 2024). To provide new insights into the functional role of intraspecies gene diversity, we have previously exposed the high-salt, high-sunlight adapted microbial communities at the end of the salt harvest cycle (∼36% NaCl) in the ‘Es Trenc’ solar salterns on the Island of Mallorca (Spain) to changing environmental conditions for about one month, and followed the communities with time-series shotgun metagenomics relative to control ponds with no treatment (i.e., ambient sunlight and salt saturation conditions) during this period (Viver 2019, Conrad 2022). The changing conditions included an experimental manipulation of light intensity through the application of a shading mesh as well as (in separate ponds) lowering salinity to ∼12% through the dilution of the brine with seawater. To aid the metagenomics, we isolated and sequenced 102 randomly-selected isolates of *Sal. ruber* from the same samples. We supplemented this genome dataset with 20 isolates recovered from a single 1-liter sample -at salt-saturation point- from the Salinas del Carmen solar salterns on the Canary Islands (>2,000 km away from Mallorca), 63 isolates from a single sample from Santa Pola’s salterns (mainland Southern Spain, 300 km away from Mallorca) as well as 9 available *Sal. ruber* genomes from the NCBI database. The ANI value patterns among all these genomes has revealed a pronounced gap around 99.5% ANI, consistent with the previous literature mentioned above and justifying *Sal. ruber* as a model system to study recombination patterns and speciation (Viver 2024).

We compared the available *Sal. ruber* isolate genomes of varied genomic relatedness to each other, ranging from members of the same genomovar (ANI >99.5%) to members of increasingly more divergent genomovars (ANI in the 97-99% range). The available genomes form six major clades (or phylogroups) based on ANI values or core-genome phylogeny, showing about 97.5-98% ANI among the phylogroups vs. 98-99.5% within a phylogroup (for members of different genomovars of a phylogroup), providing a gradient of relatedness (Fig 1). When we examined the nucleotide sequence identity patterns of individual genes across the whole genome, we observed that members of the same genomovar are typically identical or almost identical (nucleotide identity >99.8%) in most of their genes (>80% of the total, typically), except for a few regions (hotspots) that have accumulated substantial sequence diversity (e.g., showing 95-99% nucleotide identity to other members of the same genomovar; Fig. 2, top two genomes). Intriguingly, in about half of the cases, the genes in the hotspots of diversity have an identical match to another *Sal. ruber* genome of a different genomovar in our collection, indicating recent horizontal gene transfer (HGT) events mediated by homologous recombination from that genomovar or its recent ancestors (Fig. 2, blue arrows; and Fig. S1 for a phylogenetic tree-based evaluation). It is thus likely that the other half of the cases are also the product of recent HGT, but we did not have the donor genome among our isolate collection. Consistent with this interpretation, our previous study indicated that at least a couple thousand genomovars make up the natural *Sal. ruber* population in the salterns, most of which were low-abundance (rare) at the time of our sampling (Viver 2024), and here we have only sequenced representative members of about 200 of these genomovars. Most of the genes in these hotspots represented core (shared) genes of the species although several accessory (or variable) genes were also noted. Alternatively, these divergent genes (and hotspots of diversity) could represent regions of hyper-mutation, but this scenario appears less likely given that the predicted functions of the divergent genes are, more or less, random subsections of the total functions in the genome (Fig. 3) and do not show increased non-synonymous (pN) mutations (Fig. S2). Hence, the increased sequence diversity between members of the same genomovar is unlikely to represent hyper-mutation or positive (adaptive) selection. Further, the length of the (presumed) recombined segments, using the total length of consecutive recombined genes as a proxy, was similar to that observed in previous laboratory recombination studies (Power 2021) and ranged between 1 and 20 kbp, with the majority being 1-3 kbp long (Fig. S3) Therefore, it appears that these *Sal. ruber* genomes are engaging in genome-wide, rapid recombination that affects sequence identity much more than diversifying (point) mutations, revealing a recombinogenic rather than clonal sequence evolution.

**Figure 1.**
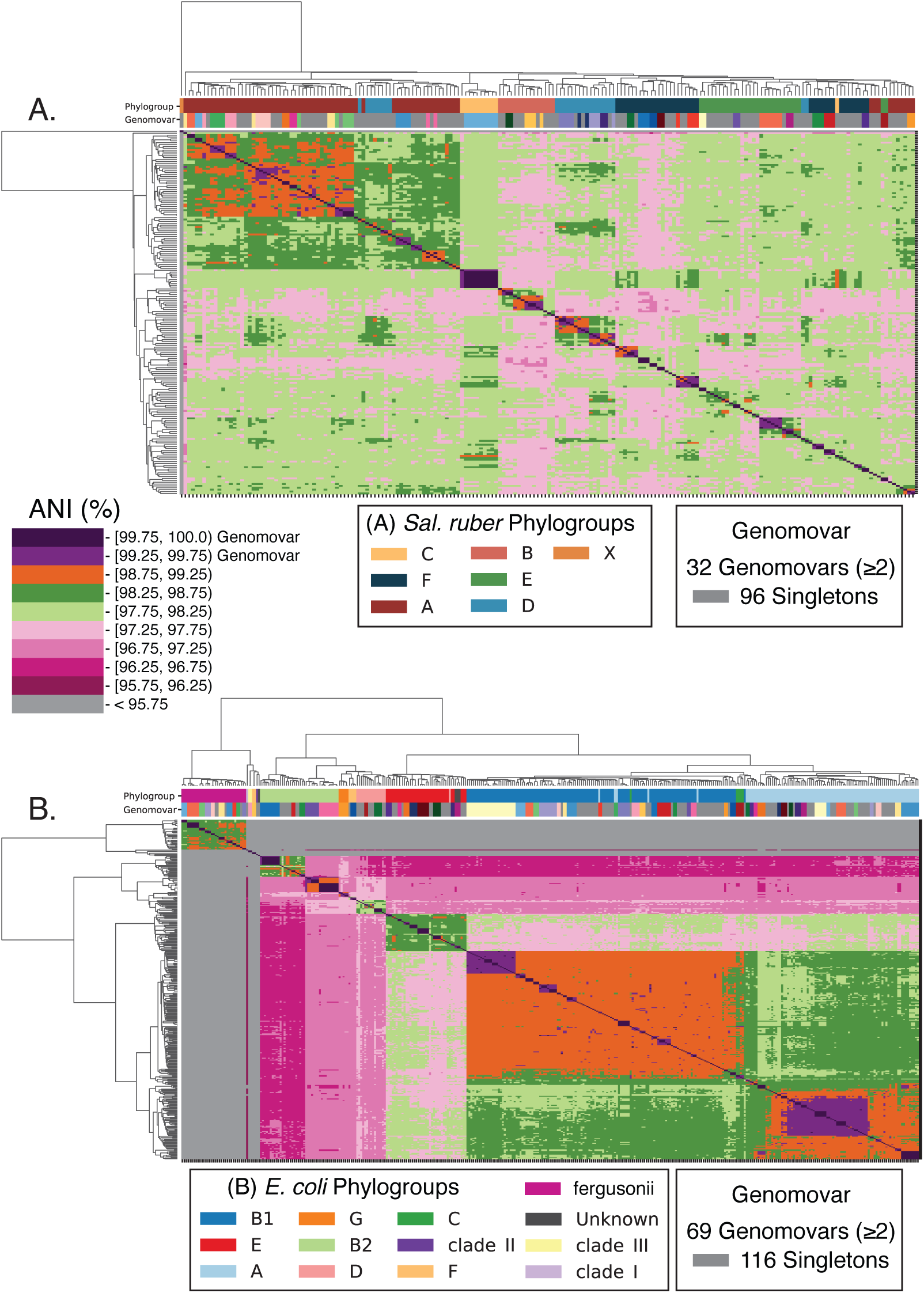
ANI clustering showing genomovar and phylogroup structure for the *Sal. ruber* and *E. coli* genomes used in this study. All vs. all ANI values were computed for *Sal. ruber* (A) and *E. coli* and relatives (*E. fergusonii / Escherichia* clades I-III) (B) using FastANI with default settings. Hierarchical clustering was performed with average linkage using Euclidean distances. Phylogroups were determined from a concatenated core gene tree for each species and with Clermon Typing. Genomovar assignments were called based on ANI values.

**Figure 2.**
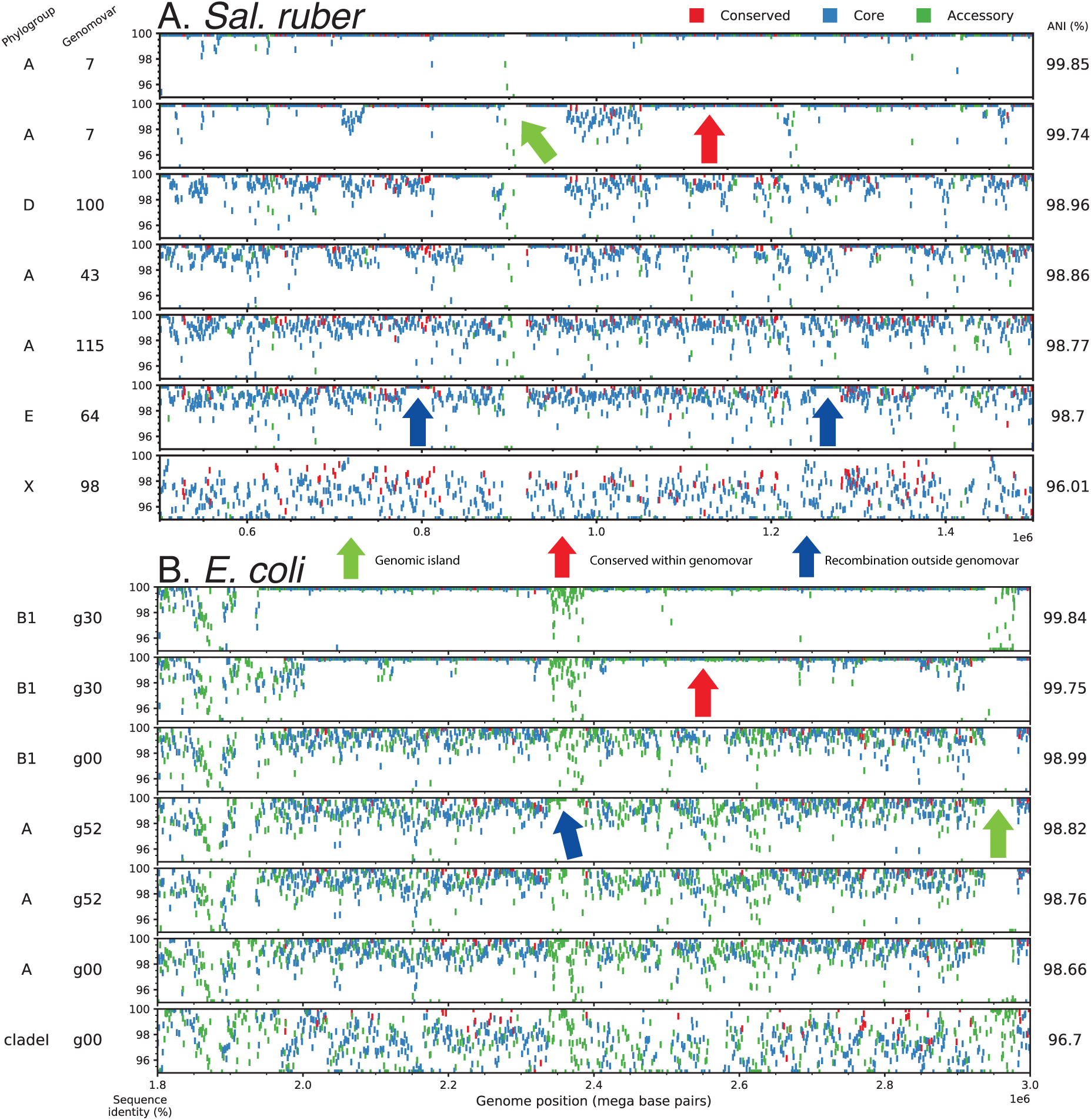
Extensive recent recombination within the *Sal. ruber* and *E. coli* genomes. Pairwise reciprocal best match (RBM) genes were identified for eight *Sal. ruber* (A) and eight *E. coli* (B) genomes spanning different genomovars and clades/phylogroups using BLAST+ with default settings. Each rectangular marker represents a gene, colored differently for highly conserved/universal, core, and accessory genes (see key), and represents the nucleotide sequence identity of RBM genes (y-axis) shared between seven query genomes (each row) and the same reference genome (x-axis, RBM gene position in reference genome) sorted by their ANI values to the reference genome shown on the far right of the panels. Two genomes from the same genomovar as the reference genome are shown in the top two rows and other genomovars and phylogroups are shown below. Note the hotspots of sequence diversity among members of the same genomovar, and that some of the genes in these hotspots show ∼100% nucleotide identity between the reference genome and genomes of other genomovars (e.g., blue arrows). Green arrows denote genomic islands specific to the reference genome (i.e., not shared with query genomes, denoted by lack of markers in the genomes not carrying the island in the corresponding region of the reference genome) while red arrows denote highly identical regions conserved within the genomovar.

**Figure 3.**
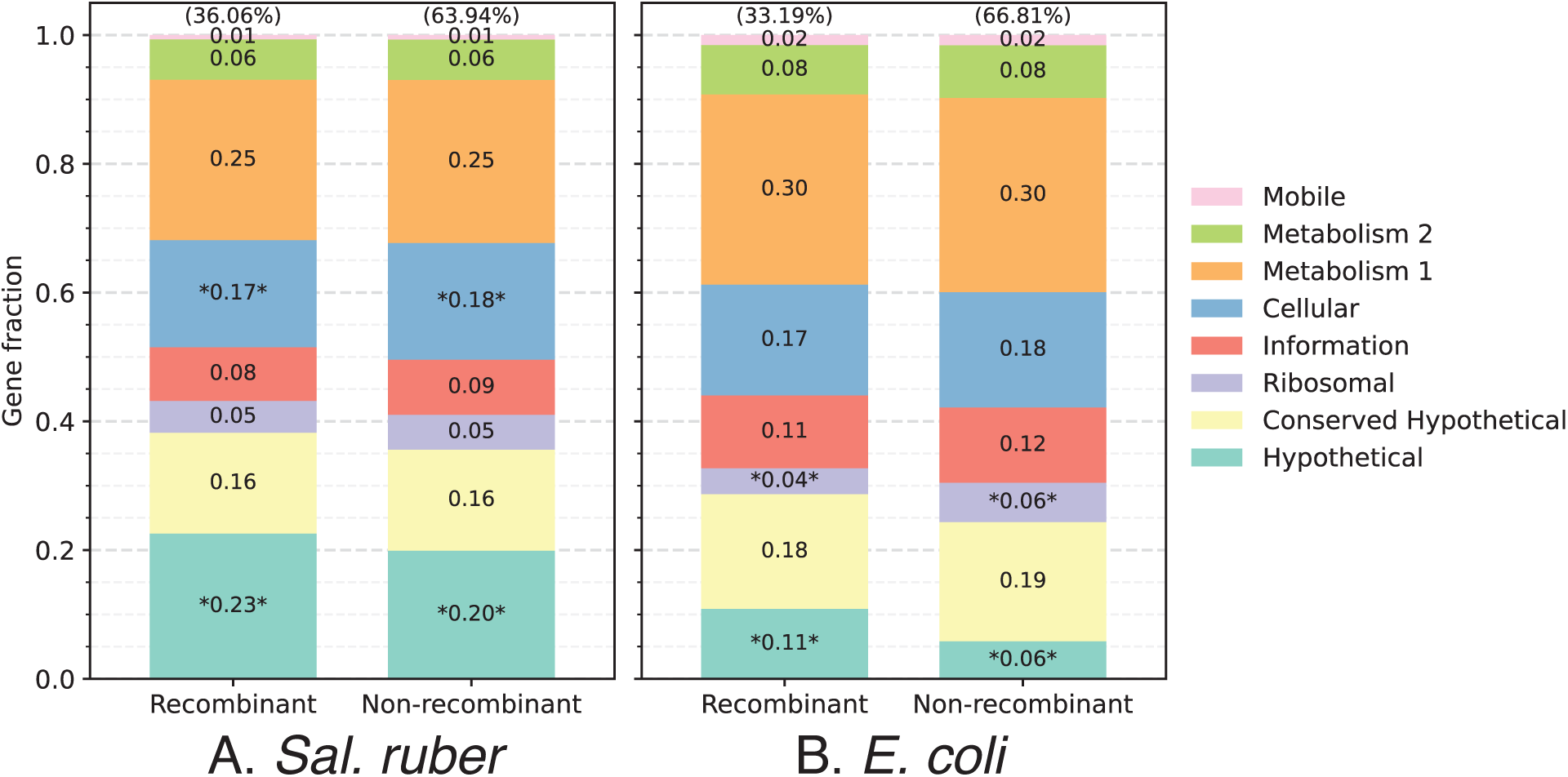
Limited functional biases in the recently recombined genes. The graphs show gene annotations summarized by high-level COG categories as a fraction of total genes in the genome (y-axis) for RBM genes divided into two categories (x-axis): genes with ≥ 99.8% sequence identity (recombinant), and genes with < 99.8% sequence identity (non-recombinant). The asterisks represent functional categories found to be significantly different by chi-square test, likely reflecting genes undergoing more frequent recombination than the average gene in the genome, favored by selection for the corresponding functions. Nonetheless, note that, overall, all functional categories are subject to recombination (left columns) and, more or less, with the same frequency -or distribution- as they are found in the genome (right columns) for both species.

To estimate more precisely the relative contribution of homologous recombination compared to diversifying mutation, we developed an empirical approach based on the sequence identity patterns across the genome. Our approach identifies recombined genes that represent recent events, i.e., showing 99.8- 100% nucleotide sequence identity, and subsequently calculates how much sequence divergence these events presumably removed based on the ANI of the genomes compared. For example, if the ANI of the two genomes compared is 97%, this would mean that the divergence of the recombined genes was about 3%, on average, before the recombination took place, and thus recombination should have removed (purged) a total of nucleotide differences that should roughly be equal to [3% × total length of recombined genes]. During the same evolutionary time, diversifying mutation can create nucleotide differences that should roughly be equal to [average divergence of recombined genes × total genome length] (because we are focusing on recent evolution only that corresponds to the 99.8-100% identity threshold used to identified recent recombination events or 0.00-0.2% sequence divergence accumulated). Using this approach, we observed that the ratio of mutations purged by homologous recombination (r) vs. mutations created by point mutation within the same time (m), or simply the r/m ratio, to be higher than 1 and often around 3-5 for several genome pairs, especially members of different genomovars of the same phylogroup (Fig. S4). In contrast, the r/m ratio between *Sal. ruber* genomes and those of *Sal. pepae*, the closest known relative that often co-occurs with *Sal. ruber* in the salterns and shares high genetic (ANI ∼94%) and metabolic relatedness (Viver 2023), was always much lower than 1 (Fig. S4). Further, genomes of different phylogroups showed typically lower r/m values than genomes of the same phylogroup, often around 1 or sometimes lower (denoted by points with ANI <98% in Fig. S4). Note that it is not feasible to perform this type of analysis for members of the same genomovar due to the high identity across the whole genome (i.e., there is no signal over the background level of sequence identity to detect recombination), and thus our r/m ratio estimates for this level (i.e., ANI >99.5%) are not reliable.

We also assessed the distribution of the identified exchanged genes across the genome to reveal whether recombination affected all regions of the genome (random distribution), and could lead to recombinogenic species and unit coherence, or if instead the exchanged genes are spatially located in a few regions across the genome (biased distribution). The latter pattern would indicate selection-driven genetic exchange (not recombinogenic speciation), and ecological speciation. Our analysis showed that while there are regions with increased frequency of recombination relative to the average of the genome, overall, no region longer than 100-200 kbp exists in the genome for which the importance of recombination is not greater than that of point mutation in at least a couple pairs of genomes from different genomovars (Fig. S5). Further, while the fraction of the genome affected by recombination between any two genomes (of different genomovars) was almost always less than 50% of the total length (pairwise comparisons), when we compared one reference genome against representative genomes of all available genomovars, this fraction often approached 80% or higher when all recombination events detected with all possible partners in the analysis were summed (Fig. 4 and S6; one vs. many comparisons). Such results were obtained with all reference genomes used in the analysis and did not appear to be specific to one or a few (reference) genomes or clades. That is, almost the whole genome was found to have recently recombined when all *Sal. ruber* genomes were considered in the analysis. Therefore, it appears that, for the *Sal. ruber* genomes evaluated here, homologous recombination is frequent and random (spatially across the genome) enough to serve as the mechanism for species cohesiveness. Further, we did not observe any strong biogeographical patterns (i.e., diversity to be locally constrained) in our recombination analysis; e.g., genomes from the Mallorca Island shared recent recombination events with genomes recovered from the Canary Islands or mainland Spain (Santa Pola) (Fig. S7). Therefore, it appears that the recombination patterns reported here apply to the global *Sal. ruber* population, not just the local populations.

**Fig 4.**
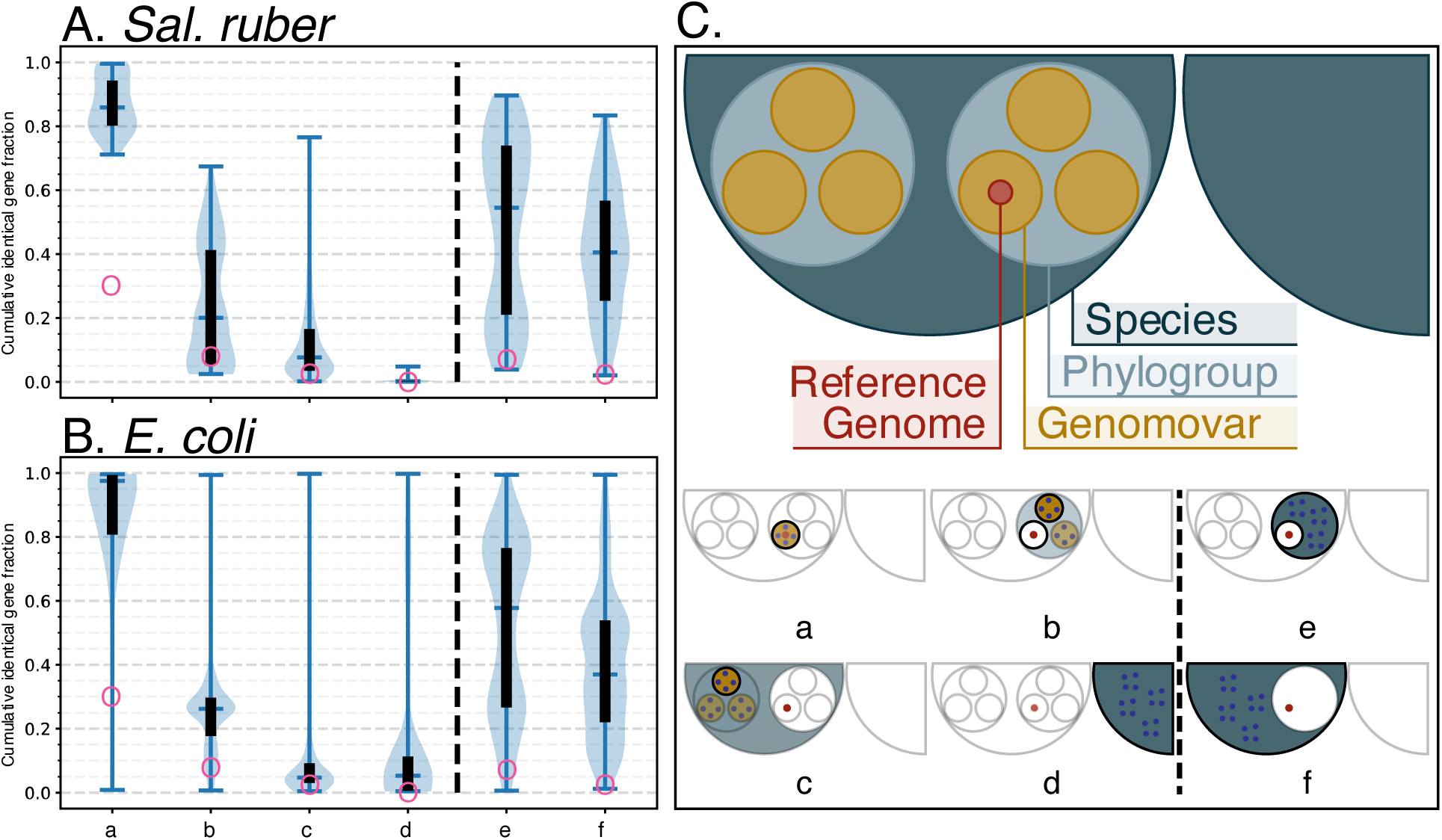
Fraction of identical genes a genome shares with all other genomes within or between genomovar, phylogroup, and species. Each genome was compared to all other genomes within each grouping (a-f) and the cumulative fraction of shared identical genes was recorded and plotted using the custom script *Allv_RBM_Violinplot.py* (i.e. each data point in the figure denotes the cumulative identical gene fraction as a function of genomes (g) where g is the total genomes in each grouping). The groupings were as follows: (a) genomes within the same genomovar, (b) genomes in each separate genomovar within the same phylogroup, excluding genomes from the same genomovar, (c) genomes in each separate genomovar within different phylogroups, (d) genomes of the other species (*S. pepae* for *Sal. ruber* and *E. fergusonii* for *E. coli*), (e) genomes within the same phylogroup excluding genomes from the same genomovar, (f) genomes within the same species excluding genomes from the same phylogroup. Panel C shows a graphical representation for comparisons performed. Data are presented in hybrid violin plots where the top and bottom whiskers show the minimum and maximum values, the middle whisker shows the median value, the black boxes show the interquartile range, and the shaded light blue regions show the density of values along the y-axis. See also Figure S6 for graphical examples of the underlying data. Note that while one or a few genomes create extreme outliers, overall, the fraction of identical genes gradually decreases among more divergent genomes compared. Also, note that our modeling analysis (red circles on the graph; see Methods for more details) suggests -for example- that only about 6-7% of the total genes in the genome should be expected to be identical among genomes showing around 98.5% ANI if there is no recent recombination (i.e., the b and e groups); both species show many more such genes in one-to-one genomovar (group B) or one-to-many genomovars (group e) at this level, revealing extensive recent gene exchange.

Our analysis also showed that genetic exchange between members of (distinct) genomovars of the same species is much more frequent than with members of different species. For instance, new gene exchange that creates genomic islands through presumably non-homologous recombination is comparatively much less frequent compared to shared-gene exchange via homologous recombination according to our comparisons among the *Sal. ruber* genomes (Figs. 2 and 4). Similarly, in the sampled salterns where *Sal. pepae* was indeed found to co-occur with *Sal. ruber* (ANI between the two species is ∼94%) (Viver 2023), our comparisons show that the two species rarely exchange shared (core) genes via a homologous recombination mechanism, at least 10 times less frequent than within species gene exchange (Fig. 4), consistent with the r/m ratio results mentioned above (Fig. S4) and the low efficiency of homologous recombination at this level of genetic relatedness. For instance, recombination efficiency drops by about 5- fold when the recombined sequences show ∼99% nucleotide identity vs. 95% identity, and by 10-fold with 90% identity (Power 2021, Konstantinidis 2023). Therefore, shared-gene exchange via homologous recombination and similar, but not necessarily fully overlapping, ecological niches, appear to keep these genomes as members of the same, sequence-discrete *Sal. ruber* species, and accounts for the 95% ANI gap at the species level.

Since recombination between members of different genomovars is quite frequent based on our evaluation (Figs. 2, 4 and S7), we hypothesize that recombination is even more frequent within members of the same genomovar, and this may account for the high identity in the rest of the genome within a genomovar. This hypothesis is also supported by the fact that homologous recombination is known to be more efficient with higher sequence similarity (Power 2021), which is the case for members of the same (ANI >99.5%) vs. different (ANI between 97-99%, typically) genomovars. Accordingly, our working model of how genomovars are maintained involves high recombination with members of the same genomovar when these members share the same ecological niche/habitat, and thus frequently encounter each other. And, this process accounts for (or leads to) the 99.5% ANI gap at the genomovar level. In contrast, when members of a genomovar are physically separated from each other, they may engage in recombination with co-occurring members of other genomovars which could then lead to the rapid emergence of new genomovars. Consistent with this model, we have evidence that at least some *Sal. ruber* genomovars show distinct ecological preferences, e.g., they prefer low -as opposed to saturation level- salt conditions (Viver 2024). Therefore, it is conceivable that members of the same genomovar grow together when growth conditions for the genomovar are favorable, and consequently there is more of a chance for recombination between them vs. with members of different genomovars that prefer different growth conditions during these periods, which eventually leads to the 99.5% ANI gap. It should be mentioned, however, that we are currently unable to directly test this hypothesis (e.g., detect recombination events between members of a genomovar) because members of the same genomovar usually share ∼100% sequence identity (no signal over the background identity level).

Alternatively, it is possible that the 99.5% ANI gap might be driven by the recent reproduction (a blooming event) of a few cells, members of the same genomovar, followed by rapid recombination of some of the offspring cells with members of other genomovars. Such inter-genomovar recombination events then lead to only a few genomes showing ANI values around 99.5% with the blooming (sub-)population of cells since inter-genomovar recombination typically involves partners sharing <99% sequence identity, causing the quick divergence of the recombining genomes from the dominant sub-population. That is, recombination with other genomovars is the dominant process, not recombination within a genomovar, and has a genome/population-diversifying rather than a population-cohesion role under this hypothesis. We are unable to directly test this alternative hypothesis for the reasons mentioned above currently (e.g., low signal over the background identity within the genomovar). However, it is intriguing to hypothesize that the same mechanism that drives the species gap, with recombination being the force of cohesion as described above, may also drive the genomovar gap, as opposed to a distinct mechanism for the latter that involves diversifying recombination. Further, while rapid and random (unbiased) diversifying recombination could theoretically provide sequence-discrete clusters similar to those observed for genomovars (Straub and Zhaxybayeva 2017), obtaining clusters with the area of inter-cluster discreteness to be centered around the exact same ANI value (i.e., 99.5%) across many different taxa [(Rodriguez 2023) and below] based on random processes seems unlikely. Hence, we favor the hypothesis that recombination as a cohesive force, coupled to high ecological cohesiveness, may be the mechanism that maintains not only the species unit but also the intraspecies units revealed here and previously (Rodriguez 2023) for the time being. Consistent with this hypothesis, we commonly observed higher gene flow between genomovars of the same phylogroup (i.e., within the same phylogroup) than between phylogroups, although there are a few phylogroups with substantial inter-phylogroup gene flow as well (Fig. 4 and S7).

Finally, it is important to note that while the genomovars might have different growth preferences as we observed previously (Viver 2024), these are likely not discrete but rather partially overlapping. For instance, we have isolated genomes that apparently prefer low-salt from salt-saturation samples and *vice-versa* (Viver 2024) and all these *Sal. ruber* genomovars can withstand salt-saturation conditions. Consequently, the 99.5% ANI gap might not always be clear or the gap may appear to be shifted to other ANI values in a few cases (Rodriguez 2023), and the gap is often not as pronounced as the 95% ANI gap that usually separates species (e.g., distinct species have less overlapping ecological niches than distinct genomovars or distinct phylogroups of the same species). Therefore, for future studies, we recommend assessing the ANI value distribution for the species of interest, and if the data indicate so, to adjust the ANI threshold to match the gap in the observed distribution.

### Applicability of the results to other species

To test how broadly these findings might apply to other bacterial species, we applied the bioinformatic framework outlined above to a set of *Escherichia coli* and *Escherichia fergusonii* genomes isolated from livestock farms and runoff in the same region (∼100 km radius) (Shaw 2021). These *E. coli* genomes showed similar genomovar and phylogroup structure to the *Sal. ruber* genomes although there was a difference with three equally dominant phylogroups (B1, A, and E) among the former genomes vs. one dominant (and five less dominant) phylogroups among the latter genomes (Fig. 1). Patterns of gene exchange and r/m ratios for the *E. coli* genomes appeared remarkably similar to those of *Sal. ruber* based on the analysis of the identical gene fraction in one vs. many genome comparisons of genomovars of the same vs. different phylogroups (Figs. 2, 4 and S4). A few qualitative differences were also observed such as that *Sal. ruber* genomes showed extreme cases of high recent gene exchange between multiple phylogroups compared to *E. coli,* which showed only a few genome pairs with similarly high gene exchange and only between phylogroups B1, B2, and A (Fig. S7). There were also a few cases of high gene flow between *E. coli* and *E. fergusonii* genomes, the closest relative sharing about ANI 93% with *E. coli*, similar to the relatedness between *Sal. ruber* and *Sal. pepae*, but these appear to involve genes that are localized in a couple specific regions of the genome and encode specific (biased) functions (selection-driven). These differences could be biological or ecologically meaningful; however, they could also be due to sampling bias (e.g., a different number of genomes is available for each phylogroup), and further research is required to investigate these differences.

## CONCLUSIONS

Collectively, our results show that recent gene exchange is both frequent and random enough across the genome to serve as the mechanism of cohesion for the sequence-discrete units of at least the two taxa studied here, *Salinibacter* and *Escherichia*. The recent gene exchange appears to be mediated by homologous recombination at the genetic/molecular level and by high ecological similarity for bringing the organisms in close, physical proximity for the genetic exchange to take place. While elements of this model have been proposed previously [e.g., (Shapiro and Polz 2015, Doolittle 2019)], it is important to note that we provide a complete mechanistic view of how the evolution of species and genomovars takes place and the necessary quantitative data in support of the model, which -to the best of our knowledge- is unprecedented. Further, instead of attributing the sequence-discrete units to either ecological or genetic (e.g., recombination) mechanisms, our model suggests that the two types of mechanisms may operate together, which represents a departure from previous models of speciation. Our results also suggest that bacteria, and likely other microbes, may evolve more “sexually” than previous thought. The drastically different lifestyles of the two taxa studied here, with *E. coli* being a human/animal gut commensal and *Sal. ruber* a halophilic environmental bacterium, as well as their large phylogenetic distance, as members of distinct bacterial phyla, indicate that the results reported here are likely applicable to additional taxa. In fact, our recent work suggests that similar patterns of recent gene exchange can be observed in human and bacterial viruses (Aldeguer et al, in review). Therefore, it is highly likely that the model of genome evolution and speciation proposed here applies more broadly in the microbial world.

## MATERIALS AND METHODS

All genomes were downloaded from NCBI’s Assembly database. Accession numbers are available in supplemental file 1. The *Salinibacter ruber* genomes reported here that were isolated from Santa Pola were sequenced as part of the GEBA V project. Step by step details for our main analysis workflow are outlined in a GitHub repository: https://github.com/rotheconrad/F100_Prok_Recombination. Briefly, ANI was calculated with FastANI v1.33 with default settings (Jain 2018). Phylogroups for *Sal. ruber* were determined from a concatenated core gene tree and hierarchical ANI tree. Phylogroups for *E. coli* were retrieved from Shaw 2021 who assigned them with ClermonTyping v1.4.1 (Beghain 2018). Genomovar assignments were called manually based on hierarchical ANI clustering. Hierarchical clustering was performed in Python 3.6+ using Seaborn v0.12.1 (Waskom 2021) and SciPy v1.9.3 (Virtanen 2020) function scipy.cluster.hierarchy.linkage with parameters method=’average’, metric=’euclidean’. Reciprocal best match genes were computed using BLAST+ v2.13.0 with default settings (Camacho 2009). Gene predictions were called using Prodigal v2.6.3 with default settings (Hyatt 2010). Gene clustering was performed with the cluster module of MMSeqs2 with settings --min-seq-id 0.90 --cov-mode 1 -c 0.5 -- cluster-mode 2 --cluster-reassign (Steinegger and Soding 2017). Recombined genes and the ratio of recombination-to-mutation were determined as described in the main text.

Our modeling analysis to estimate the fraction of identical genes expected between two genomes of a given ANI value without any recent recombination between the genomes (i.e., red circles in Figure 4) used the following approach. A random ancestral genome sequence was generated with 3000 genes of variable length (selected from a normal distribution; mu=1000bp, stdev=250) with 10bp-long random sequence inserted between any two genes. Subsequently, daughter genomes were generated by the addition of random, single-nucleotide mutations to ancestral genes to match a gamma distribution for RBM gene sequence identity with the distribution mean fit to the desired ANI of the genome pair. We generated 10 daughter genomes from the ancestral genome for each ANI value in the range of 95-100% ANI with a step size of 0.01 for a total simulated population size of 5001 genomes. The ANI and identical gene fractions for our simulated population can be found in supplemental file 2. The resulting frequency values matched well the average frequency of such identical genes found between randomly drawn genomes from NCBI of similar ANI, which are expected to not show extensive recombination, indicating that our population genome simulation was robust. The code used to simulate these genomes is available at: https://github.com/rotheconrad/Population-Genome-Simulator.

## ACKNOWLEDGEMENTS

The work (proposal DOI: 10.46936/10.25585/60001079) conducted by the U.S. Department of Energy Joint Genome Institute (https://ror.org/04xm1d337), a DOE Office of Science User Facility, is supported by the Office of Science of the U.S. Department of Energy operated under Contract No. DE-AC02-05CH11231.

The authors would especially like to thank the whole team at Salinas d’Es Trenc and Gusto Mundial Balearides, S.L. (Flor de Sal d’Es Trenc) and of the Salinas de Fuerteventura (Salinas del Carmen) for allowing access to their facilities and their support in performing the experiments. The research at the IMEDEA was funded by the Ministry of Science and Innovation projects PGC2018-096956-B-C41 and PID2021-126114NB-C42, both also supported -in part- by European Regional Development Fund (FEDER) funds, and through the “Maria de Maeztu Centre of Excellence” accreditation to IMEDEA (CSIC-UIB) (CEX2021-001198). KTK’s research was supported, in part, by the U.S. National Science Foundation (Award No. 1831582 and No. 2129823). RRM acknowledges the financial support of the sabbatical stay at Georgia Tech supported by the grant PRX18/00048 of the Ministry of Science and Innovation. TV acknowledges the “Margarita Salas” postdoctoral grant, funded by the Spanish Ministry of Universities, within the framework of Recovery, Transformation and Resilience Plan, and funded by the European Union (NextGenerationEU), with the participation of the University of Balearic Islands (UIB). TV and RA acknowledge support by the Max Planck Society.

## AUTHOR CONTRIBUTIONS

Conceptualization: KTK, RRM, LMR, REC

Data curation: REC, CEB, TV, SNV

Formal Analysis: REC, CEB

Visualization: REC, CEB

Funding acquisition: KTK, RRM, RA

Investigation: REC, CEB, LMR, TV, BAR

Methodology: REC, CEB, LMR, TV, BAR

Project Administration: KTK, RRM

Supervision: KTK, RRM, RA, SNV

Writing – original draft: KTK, REC, CEB, BAR

Writing – review & editing: REC, CEB, TV, LMR, BAR, JKH, SNV, RA, RRM, KTK

## DATA AND MATERIALS AVAILABILITY

All data are available in the main text or the supplementary materials.

## Materials and Methods

### Tanglegrams (Figure S1)

Regions of between-phylogroup recombination were identified using the *Sal. ruber* graph in Figure 2A. We selected core genes from these regions (∼800 000 bp and ∼1 250 000 bp) and extracted the gene sequences from the Prodigal files using seqTK v1.3-r117 (Li, 2013). Sequences were aligned with MUSCLE v3.8.31 (Edgar, 2004) and evaluated to ensure they were of good quality. MEGA11 (Tamura, 2021) was used to generate individual maximum likelihood trees that were compared to the *Sal. ruber* ANI cladogram shown in Figure 1 and S1. Tanglegrams were drawn in R using the ape v5.7.1 (Paradis, 2019), dendextend v1.17.1 (Galili, 2015) and phylogram v2.1.0 (Wilkinson, Davy, 2018) packages.

### pN vs pS (Figure S2)

The reference genome for Figure 2A was used to calculate the difference between synonymous and non-synonymous substitutions to measure the effect of selection on the genes. The calculations involved 13 genes spread across the genome at intervals of ∼250 000 bp and 20 genes from the same regions at intervals of 50 000 bp. Gene sequences were extracted from Prodigal files using seqTK (Li, 2013) and aligned by codons using the Clustal (Larkin, *et al*., 2007) module built in to MEGA11 (Tamura, 2021). The Kumar model (Nei and Kumar, 2000) was used to calculate overall mean distances for the 13 genes spread across the genome and pairwise distances for the genes in the region of interest as implemented in the script developed by T. Zhu, available at https://github.com/zhutao1009/dnds.git. The plot was generated in Python.

### ANI trees (Figure S7)

All-vs-All ANI tests were done with FastANI (Jain, *et al*., 2019) for *S. ruber* and *E. coli* independently. This data was used to construct ANI-based cladograms in R using the Euclidean distance and average cluster methods. Ape v5.7.1 (Paradis, 2019) was used to convert the cladograms into Newick-formatted trees that could be upload to iTol (Lutenic, *et al*, 2021) for further annotation. Phylogroups were annotated according to the major monophyletic clades that could be identified in the ANI tree (for *S. ruber*) and according to the groups previously identified in literature (for *E. coli*). F100 scores generated by the pipeline were used to draw connection arcs between pairs of genomes as a representation of the frequency of recombination between two genomes.

**Code for supplementary figures can be found at:** https://github.com/catbrink/Explaining-ANI-gaps-Code-for-supplementary-figures.git

---

**Figure S1:**
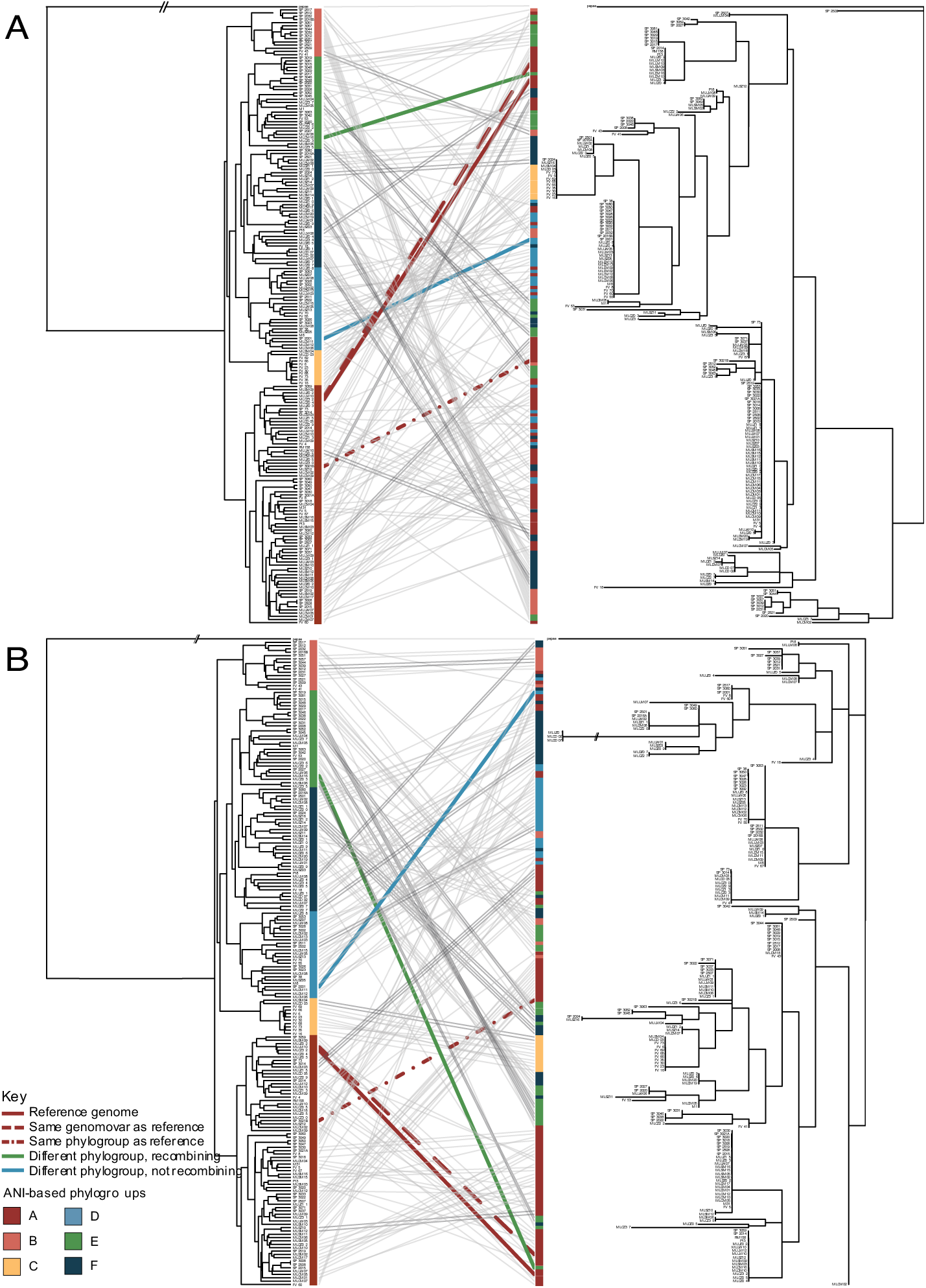
Phylogenetic trees showing examples of recent recombination of functional genes between *Sal. ruber* phylogroups. Two tanglegrams are used to illustrate the recombination in the regions identified in Figure 2A as “recombination outside genomovar” by the blue arrows. The ANI tree (left, Panels A and B) is compared to two gene trees: one from a region at 782 574 bp (right, Panel A; an adenine deaminase; closest match: UniRef100_Q2S006) and the second from the region identified at 1 261 301 bp (right, Panel B; an ATPase beta subunit or *atpD*; closest match: UniRef100_Q2RZV3) along the reference genome. The genes have undergone recent gene exchange between distinct ANI genomovars (of different phylogroups) as reflected by the clustering, at high nucleotide identity, with different genomovars in the gene relative to the ANI tree (a few examples are denoted by the colored lines). A single gene was selected from each region to generate the alignment and gene tree using MEGA11 (Tamura 2021). The base tanglegrams were constructed in the R programming language. Some branches have been shortened for display purposes, and these are indicated by two dashed lines. Phylogroups are colored as in the ANI tree.

**Figure S2:**
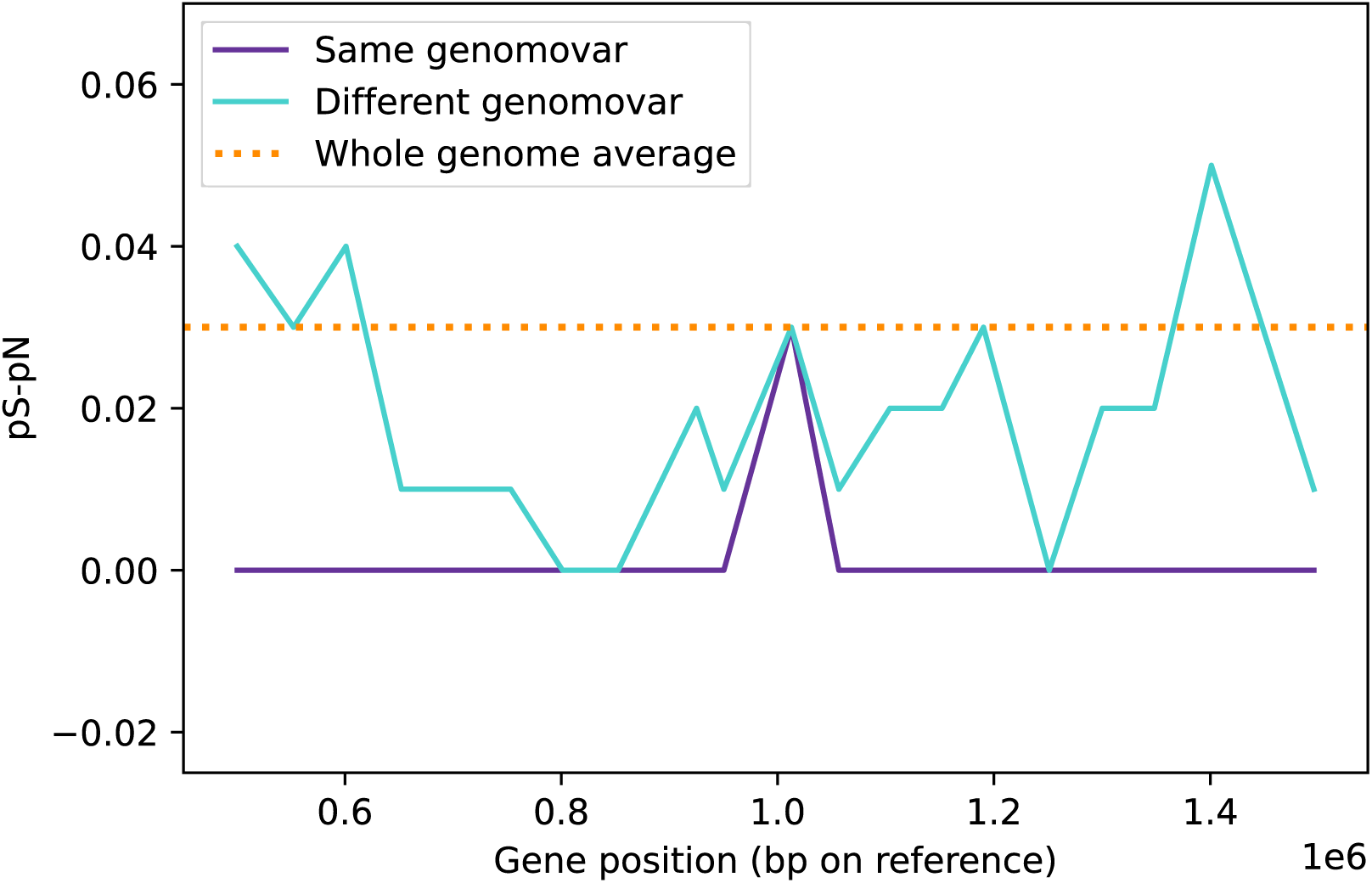
Evidence for the lack of positive (adaptive) selection in areas of recent recombination. The plot shows the ratio between the proportion of non-synonymous and synonymous substitutions along the genomic region shown in Figure 2A. The ratio fluctuates between different genomovars, but is never greater than 0.4, which indicates that it is unlikely that any of the corresponding genes are experiencing adaptive evolution. The pN/pS ratio was calculated using the script developed by T. Zhu, 2020 available at https://github.com/zhutao1009/dnds.git. The whole genome average was calculated using 13 genes spread across the whole genome at intervals of ∼250000 bp and are supplemented by ratios calculated for the same and different genomovar for 20 genes at ∼50000 bp across the segment shown in Figure 2A.

**Figure S3:**
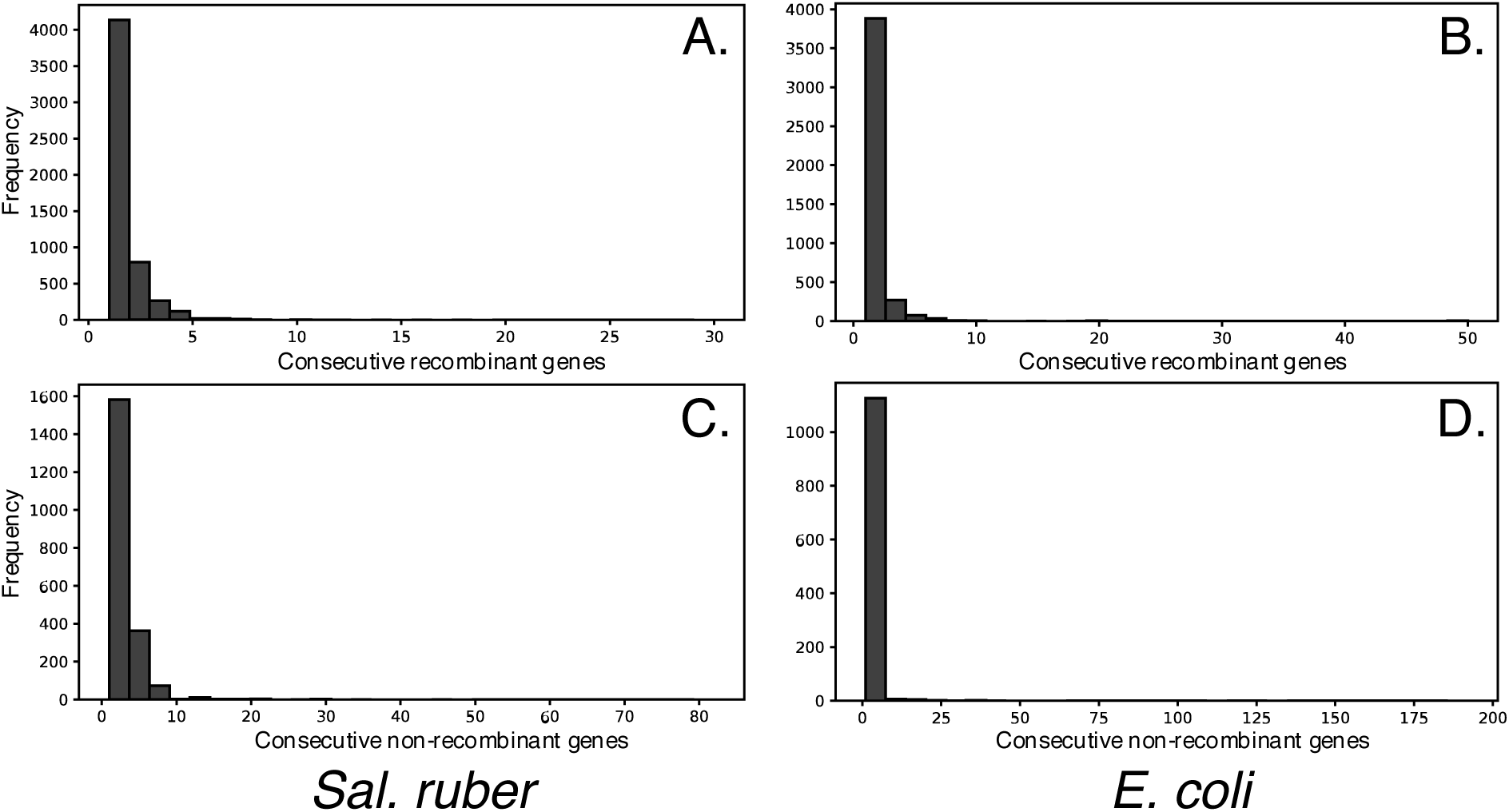
Length distribution of recombinant fragments between the genomes. The graphs show the frequency and distribution of consecutive recombinant genes as a proxy for the length distribution of recombinant fragments (A and B) and consecutive non-recombinant genes (C and D) for the *Sal. ruber* (A and C) and *E. coli* (B and D) genomes used in this study.

**Figure S4.**
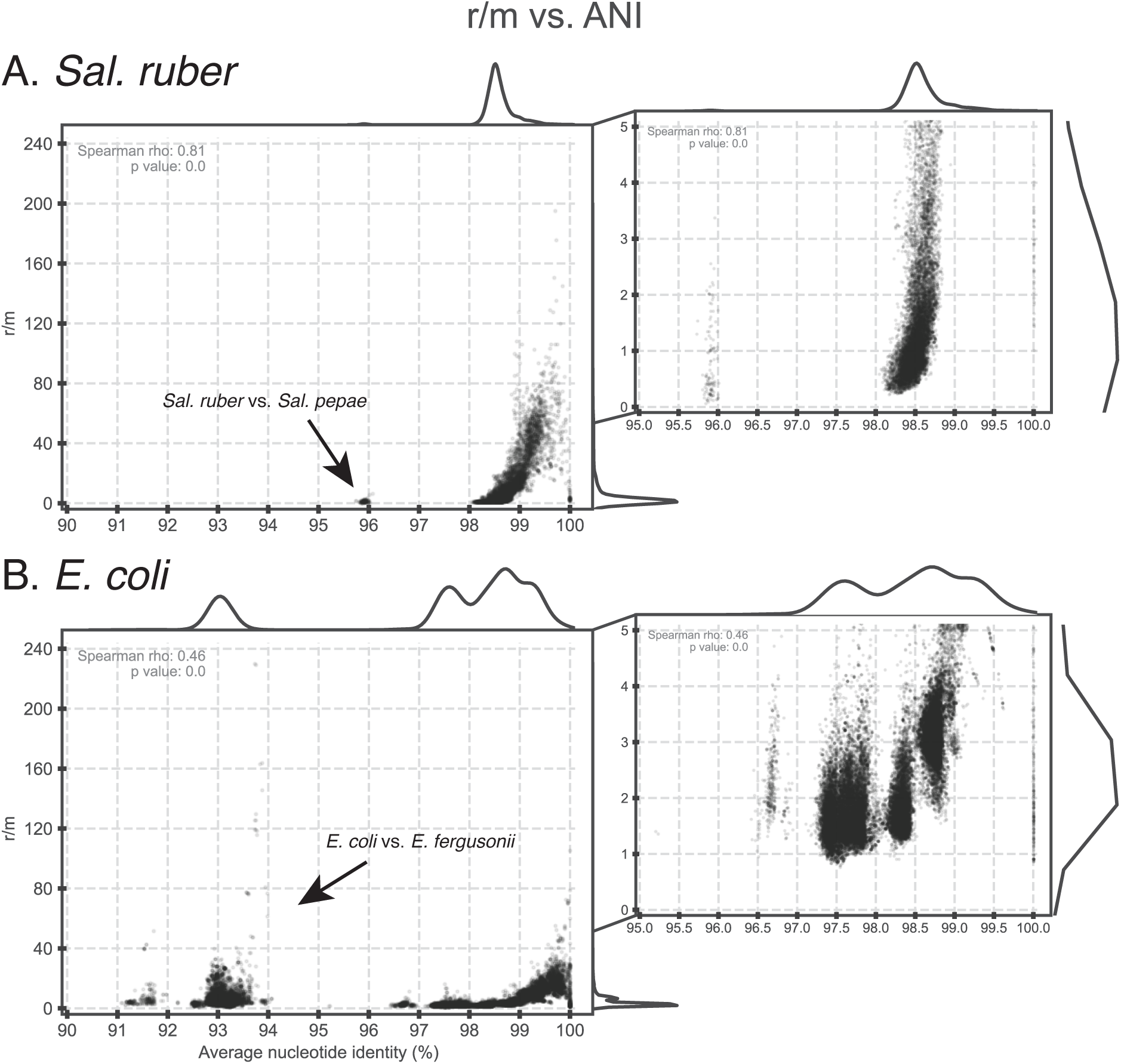
Recombination to mutation (r/m) ratio as a function of the ANI of the genome pairs compared. The r/m ratio (y-axes) was estimated for all genome pairs in our collection for each species (graph title on top) using the empirical approach described in the main text, and is plotted against the ANI value of the genome pair compared (x-axes). The marginal plots outside the two axes show histograms for the density of datapoints on each axis. Graphs on the right are zoomed-in versions of the main graphs on the left. Note that the ratio is frequently above 1 for genomes sharing between 98.5-99.5% ANI (e.g., members of different genomovars of the same phylogroup) for both species and that the estimates above ∼99.5% ANI are not reliable due to inability to detect recombination at this high sequence identity level. A few outlier datapoints (genome pairs) with ratios higher than 100 were also observed in the 98-99.5% ANI range and are due to the high identity of the recombined genes identified (causing the denominator in the r/m ratio to be a small number); the graphs on the right show the majority of datapoints, and thus better represent the average pattern. Also note that a few *E. coli* and *E. fergusoni* genome pairs (left part of the lower graph) show a ratio higher than 1, but this is driven by recombined genes that are localized in a couple specific regions of the genome and encode specific functions (selection-driven recombination, and not widespread across the genome). See main text for additional details.

**Figure S5.**
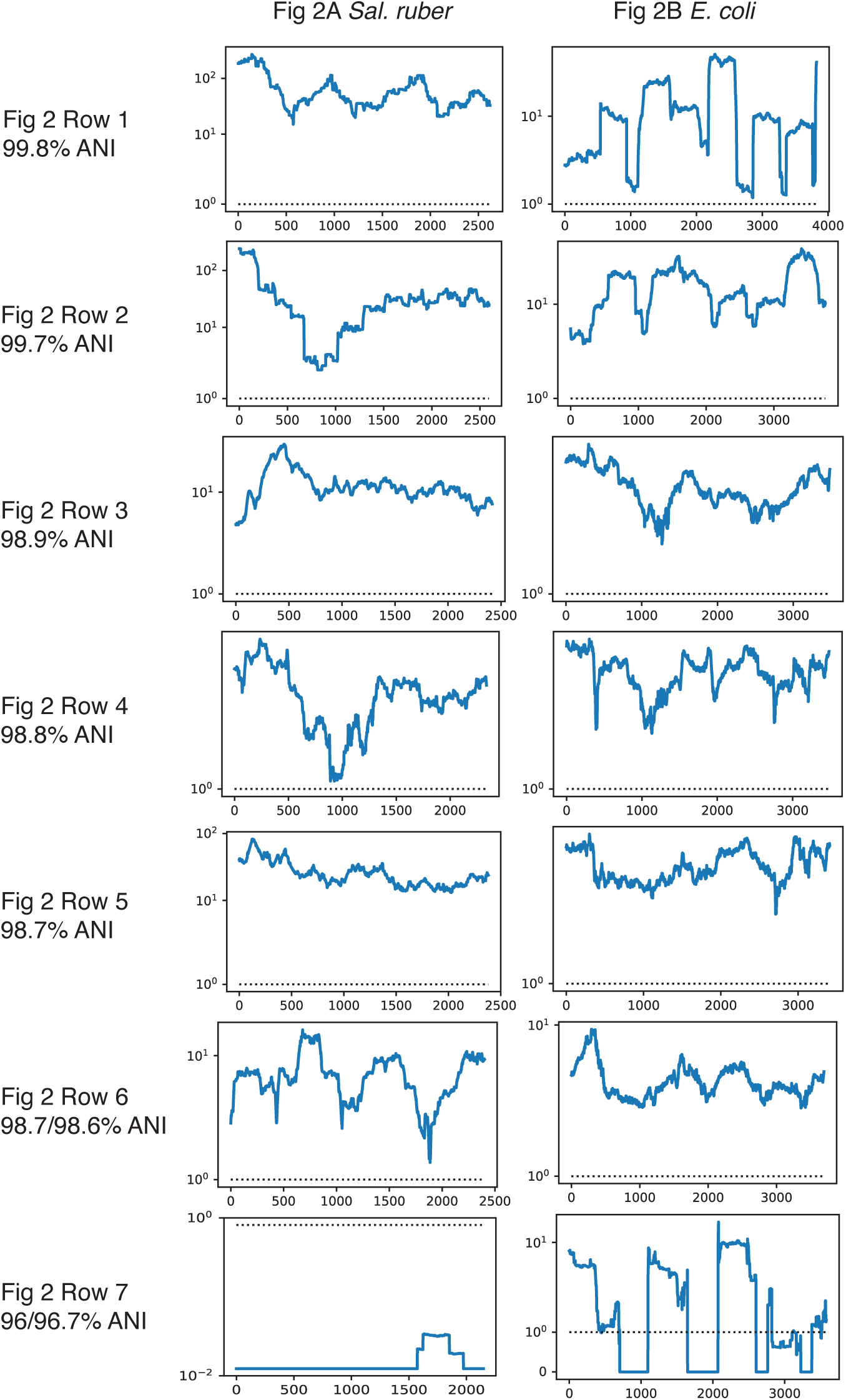
Recombination affects every segment of the genome and is higher than mutation. The same reference and query genomes, with the same order, as shown in Figure 2 were used. The graphs show the recombination to mutation ratio (r/m, y-axes) across the reference genome in pairwise genome comparisons in a sliding window of 400 genes). The ratio was estimated using our empirical approach described in the main text. Note that the ratio is often larger than 1, and even if there are segments of the genome with ratio <1 in some genome comparisons, other genomes show ratios >1 for the same segments, revealing that recombination is genome-wide (not spatially biased) at the whole-population/species level.

**Figure S6.**
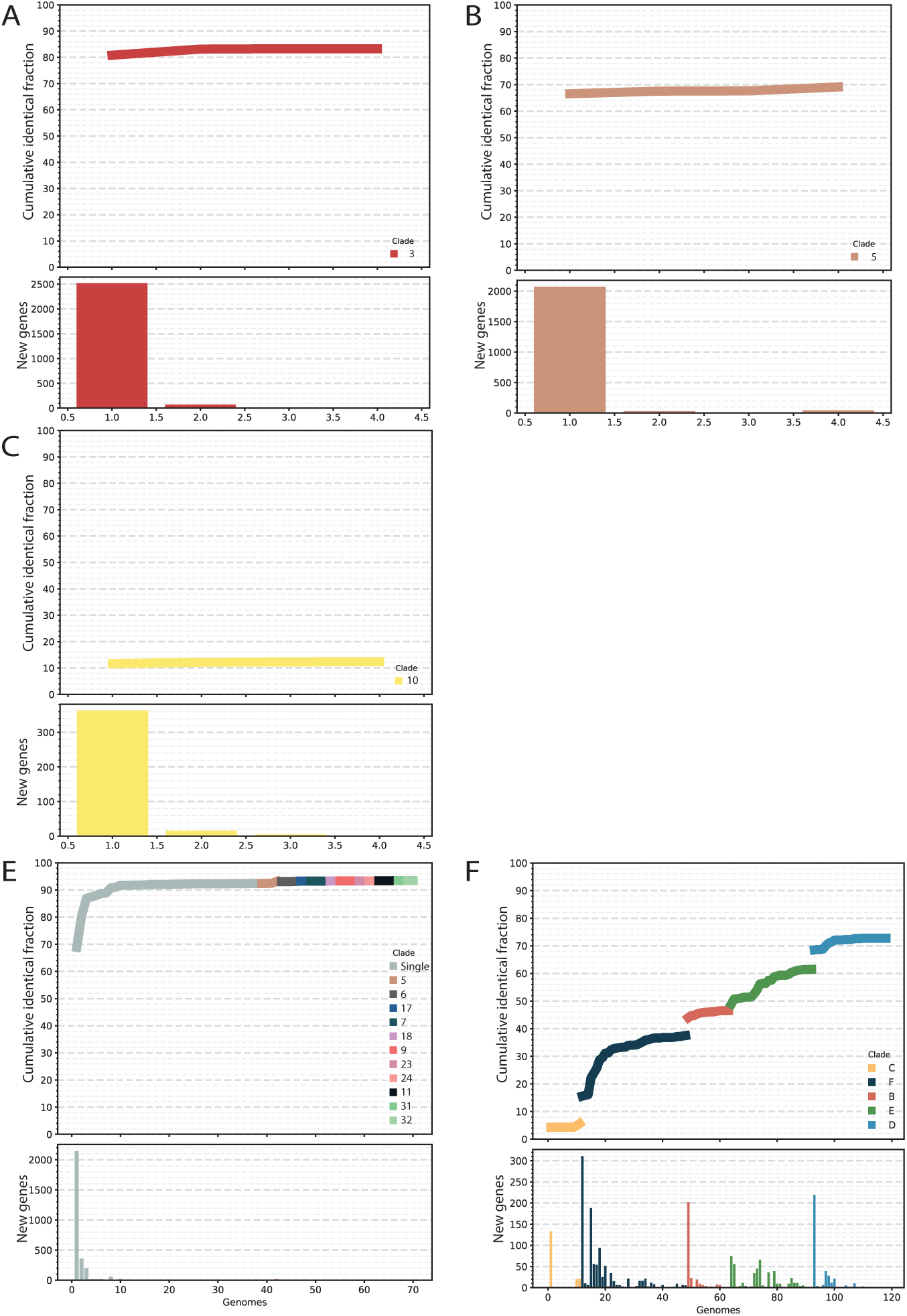
Examples of cumulative identical gene fraction curves for one vs. many genome comparisons. The y-axis shows the fraction of identical genes among all genes in the reference genome (top panels) and the number of new identical genes (bottom panels) for each genome added (x-axis), i.e. number of identical genes as a function of the number of genome partners. The same *Sal. ruber* reference genome used in Figure 4A (Phylogroup A, genomovar 3) is compared against the groupings used in Figure 4 which were (A/a) genomes within the same genomovar (genomovar 3), (B/b) genomes from a different genomovar (genomovar 5) within the same phylogroup (phylogroup A), (C/c) genomes in a different genomovar (genomovar 10) from a different phylogroup, (E/e) genomes within the same phylogroup (phylogroup A) excluding genomes from the same genomovar (genomovar 3), and (F/f) genomes within the same species excluding genomes from the same phylogroup (phylogroup A). An example corresponding to panel D (i.e., one genome against genomes of a different species) is not shown because only a single *Sal. pepae* genome was available. Figure 4 essentially shows the final (far right) point of the curve for each one vs. many genome comparison and the abovementioned groups (a though e).

**Figure S7.**
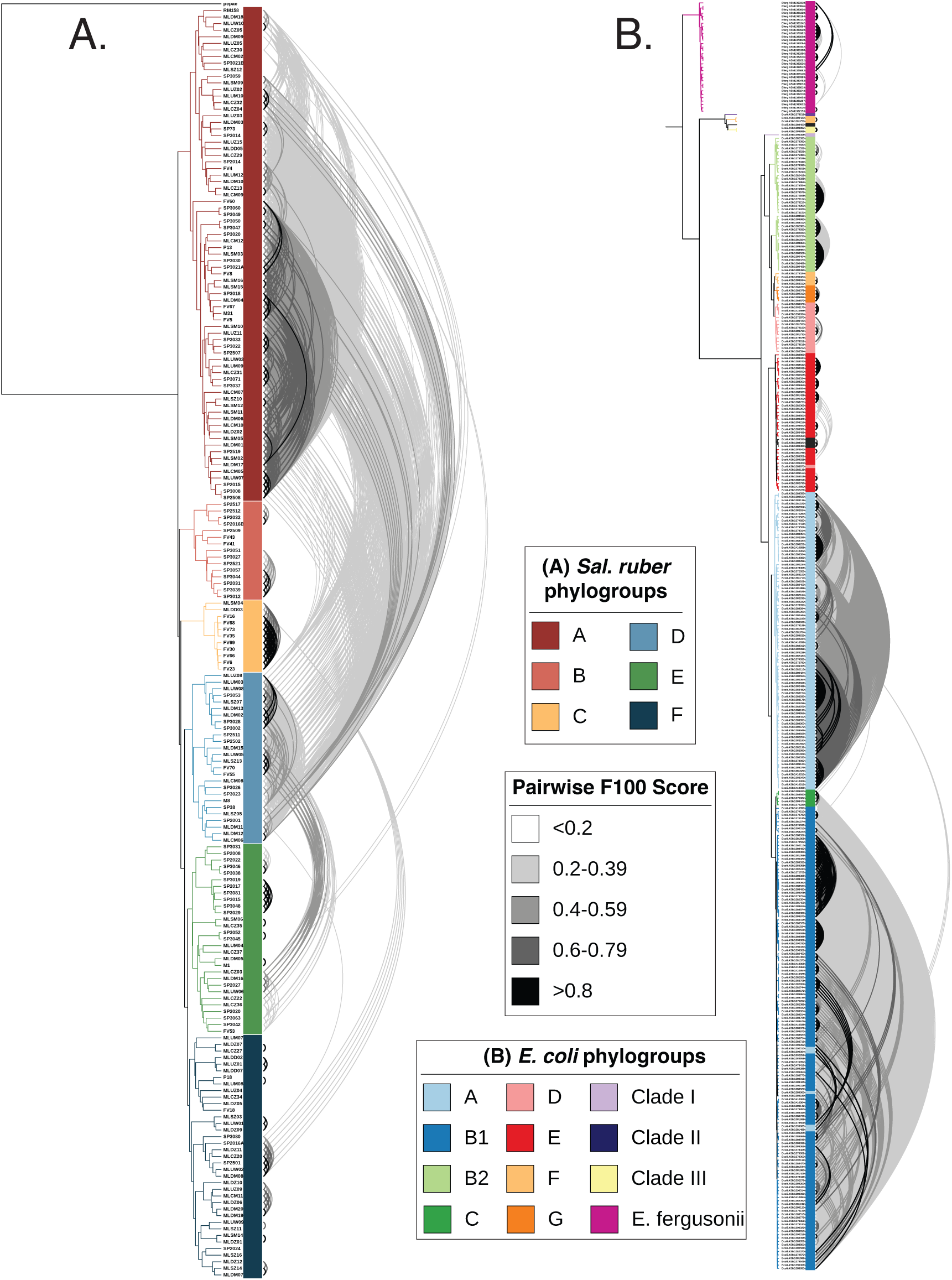
ANI trees showing the phylogroup (clade) structure for each species studied and recent gene exchange. The frequency of 100% identical RBM genes to total RBM genes found (F100), a proxy for recent recombination, for each genome pair is shown by connecting lines shaded along a grayscale gradient for *Sal. ruber* (A) and *E. coli* (B). Genome pairs with F100 value close to zero (or zero) are not shown. For *Sal. ruber*, if the name of a genome starts with ML, FV or SP, it denotes that the corresponding isolate originated from the Mallorca, Canary Islands or Santa Pola (mainland Spain) sites.

## References cited

Anton, J., R. Rossello-Mora, F. Rodriguez-Valera and R. Amann (2000). “Extremely halophilic bacteria in crystallizer ponds from solar salterns.” Appl Environ Microbiol 66(7): 3052–3057.

Beghain, J., A. Bridier-Nahmias, H. Le Nagard, E. Denamur and O. Clermont (2018). “ClermonTyping: an easy-to-use and accurate in silico method for Escherichia genus strain phylotyping.” Microb Genom 4(7).

Bendall, M. L., S. L. Stevens, L. K. Chan, S. Malfatti, P. Schwientek, J. Tremblay, … R. R. Malmstrom (2016). “Genome-wide selective sweeps and gene-specific sweeps in natural bacterial populations.” ISME J 10(7): 1589–1601.

Bobay, L. M. and H. Ochman (2017). “Biological species are universal across Life’s domains.” Genome Biol Evol.

Camacho, C., G. Coulouris, V. Avagyan, N. Ma, J. Papadopoulos, K. Bealer and T. L. Madden (2009). “BLAST+: architecture and applications.” BMC Bioinformatics 10: 421.

Caro-Quintero, A. and K. T. Konstantinidis (2012). “Bacterial species may exist, metagenomics reveal.” Environmental Microbiology 14(2): 347–355.

Caro-Quintero, A., G. P. Rodriguez-Castano and K. T. Konstantinidis (2009). “Genomic insights into the convergence and pathogenicity factors of *Campylobacter jejuni* and *Campylobacter coli* species.” J Bacteriol.

Casamayor, E. O., R. Massana, S. Benlloch, L. Ovreas, B. Diez, V. J. Goddard, … C. Pedros-Alio (2002). “Changes in archaeal, bacterial and eukaryal assemblages along a salinity gradient by comparison of genetic fingerprinting methods in a multipond solar saltern.” Environ Microbiol 4(6): 338–348.

Chase, A. B., C. Weihe and J. B. H. Martiny (2021). “Adaptive differentiation and rapid evolution of a soil bacterium along a climate gradient.” Proc Natl Acad Sci U S A 118(18).

Conrad, R. E., T. Viver, J. F. Gago, J. K. Hatt, S. N. Venter, R. Rossello-Mora and K. T. Konstantinidis (2022). “Toward quantifying the adaptive role of bacterial pangenomes during environmental perturbations.” ISME J 16(5): 1222–1234.

Doolittle, W. F. (2019). “Speciation without Species: A Final Word.” Philosophy, Theory, and Practice in Biology 11.

Drake, J. W., B. Charlesworth, D. Charlesworth and J. F. Crow (1998). “Rates of spontaneous mutation.” Genetics 148(4): 1667–1686.

Fraser, C., W. P. Hanage and B. G. Spratt (2007). “Recombination and the nature of bacterial speciation.” Science 315(5811): 476–480.

Gevers, D., F. M. Cohan, J. G. Lawrence, B. G. Spratt, T. Coenye, E. J. Feil, … J. Swings (2005). “Opinion: Re-evaluating prokaryotic species.” Nat Rev Microbiol 3(9): 733–739.

Gibbons, R. J., and B. Kapsimalis (1967). “Estimates of the overall rate of growth of the intestinal microflora of hamsters, guinea pigs, and mice.” Journal of Bacteriology 93(1): 510–512.

Gomariz, M., M. Martinez-Garcia, F. Santos, F. Rodriguez, S. Capella-Gutierrez, T. Gabaldon, … J. Anton (2015). “From community approaches to single-cell genomics: the discovery of ubiquitous hyperhalophilic Bacteroidetes generalists.” ISME J 9(1): 16–31.

Hyatt, D., G. L. Chen, P. F. Locascio, M. L. Land, F. W. Larimer and L. J. Hauser (2010). “Prodigal: prokaryotic gene recognition and translation initiation site identification.” BMC Bioinformatics 11: 119.

Jain, C., R. L. Rodriguez, A. M. Phillippy, K. T. Konstantinidis and S. Aluru (2018). “High throughput ANI analysis of 90K prokaryotic genomes reveals clear species boundaries.” Nat Commun 9(1): 5114.

Konstantinidis, K. T. (2023). “Sequence-discrete species for Prokaryotes and other microbes: A historical perspective and pending issues.” mLife: In press.

Konstantinidis, K. T., A. Ramette and J. M. Tiedje (2006). “The bacterial species definition in the genomic era.” Philos Trans R Soc Lond B Biol Sci 361(1475): 1929–1940.

Mora-Ruiz, M. D. R., A. Cifuentes, F. Font-Verdera, C. Perez-Fernandez, M. E. Farias, B. Gonzalez, …R. Rossello-Mora (2018). “Biogeographical patterns of bacterial and archaeal communities from distant hypersaline environments.” Syst Appl Microbiol 41(2): 139–150.

Murray, C. S., Y. Gao and M. Wu (2021). “Re-evaluating the evidence for a universal genetic boundary among microbial species.” Nat Commun 12(1): 4059.

Olm, M. R., A. Crits-Christoph, S. Diamond, A. Lavy, P. B. Matheus Carnevali and J. F. Banfield (2020). “Consistent Metagenome-Derived Metrics Verify and Delineate Bacterial Species Boundaries.” mSystems 5(1).

Power, J. J., F. Pinheiro, S. Pompei, V. Kovacova, M. Yuksel, I. Rathmann, … B. Maier (2021). “Adaptive evolution of hybrid bacteria by horizontal gene transfer.” Proc Natl Acad Sci U S A 118(10).

Rodriguez, R. L., R. E. Conrad, T. Viver, D. J. Feistel, B. G. Lindner, S. N. Venter, … K. T. Konstantinidis (2023). “An ANI gap within bacterial species that advances the definitions of intra-species units.” mBio: e0269623.

Rodriguez, R. L., C. Jain, R. E. Conrad, S. Aluru and K. T. Konstantinidis (2021). "Reply to: "Re-evaluating the evidence for a universal genetic boundary among microbial species"." Nat Commun 12(1): 4060.

Rossello-Mora, R. and R. Amann (2015). “Past and future species definitions for Bacteria and Archaea.” Syst Appl Microbiol 38(4): 209–216.

Roux, S., D. Paez-Espino, I. A. Chen, K. Palaniappan, A. Ratner, K. Chu, … N. C. Kyrpides (2021). “IMG/VR v3: an integrated ecological and evolutionary framework for interrogating genomes of uncultivated viruses.” Nucleic Acids Res 49(D1): D764–D775.

Seabolt, M. H., K. T. Konstantinidis and D. M. Roellig (2021). “Hidden Diversity within Common Protozoan Parasites as Revealed by a Novel Genomotyping Scheme.” Appl Environ Microbiol 87(6).

Shapiro, B. J. and M. F. Polz (2015). “Microbial Speciation.” Cold Spring Harb Perspect Biol 7(10): a018143.

Shaw, L. P., K. K. Chau, J. Kavanagh, M. AbuOun, E. Stubberfield, H. S. Gweon, … R. consortium (2021). “Niche and local geography shape the pangenome of wastewater- and livestock-associated Enterobacteriaceae.” Sci Adv 7(15).

Sheppard, S. K., N. D. McCarthy, D. Falush and M. C. Maiden (2008). “Convergence of Campylobacter species: implications for bacterial evolution.” Science 320(5873): 237–239.

Simmonds, P., M. J. Adams, M. Benko, M. Breitbart, J. R. Brister, E. B. Carstens, … F. M. Zerbini (2017). “Consensus statement: Virus taxonomy in the age of metagenomics.” Nat Rev Microbiol 15(3): 161–168.

Steinegger, M. and J. Soding (2017). “MMseqs2 enables sensitive protein sequence searching for the analysis of massive data sets.” Nat Biotechnol 35(11): 1026–1028.

Strachan, C. R., X. A. Yu, V. Neubauer, A. J. Mueller, M. Wagner, Q. Zebeli, … M. F. Polz (2023). “Differential carbon utilization enables co-existence of recently speciated Campylobacteraceae in the cow rumen epithelial microbiome.” Nat Microbiol 8(2): 309–320.

Straub, T. J. and O. Zhaxybayeva (2017). “A null model for microbial diversification.” Proc Natl Acad Sci U S A 114(27): E5414–E5423.

Virtanen, P., R. Gommers, T. E. Oliphant, M. Haberland, T. Reddy, D. Cournapeau, … C. SciPy (2020). “SciPy 1.0: fundamental algorithms for scientific computing in Python.” Nat Methods 17(3): 261–272.

Viver, T., R. E. Conrad, M. Lucio, M. Harir, M. Urdiain, J. F. Gago, … R. Rossello-Mora (2023). “Description of two cultivated and two uncultivated new Salinibacter species, one named following the rules of the bacteriological code: Salinibacter grassmerensis sp. nov.; and three named following the rules of the SeqCode: Salinibacter pepae sp. nov., Salinibacter abyssi sp. nov., and Salinibacter pampae sp. nov.” Syst Appl Microbiol 46(3): 126416.

Viver, T., R. E. Conrad, L. H. Orellana, M. Urdiain, J. E. Gonzalez-Pastor, J. K. Hatt, … R. Rossello-Mora (2020). “Distinct ecotypes within a natural haloarchaeal population enable adaptation to changing environmental conditions without causing population sweeps.” ISME J.

Viver, T., R. E. Conrad, R. L. Rodriguez, A. S. Ramirez, S. N. Venter, J. Rocha-Cardenas, … R. Rossello-Mora (2024). “Towards estimating the number of strains that make up a natural bacterial population.” Nat Commun 15(1): 544.

Viver, T., L. Orellana, P. Gonzalez-Torres, S. Diaz, M. Urdiain, M. E. Farias, … R. Rossello-Mora (2018). “Genomic comparison between members of the Salinibacteraceae family, and description of a new species of Salinibacter (Salinibacter altiplanensis sp. nov.) isolated from high altitude hypersaline environments of the Argentinian Altiplano.” Syst Appl Microbiol 41(3): 198–212.

Viver, T., L. H. Orellana, S. Diaz, M. Urdiain, M. D. Ramos-Barbero, J. E. Gonzalez-Pastor, … R. Rossello-Mora (2019). “Predominance of deterministic microbial community dynamics in salterns exposed to different light intensities.” Environ Microbiol 21(11): 4300–4315.

Waskom, M. L. (2021). “Seaborn: statistical data visualization.” Journal of Open Source Software 6: 3021.

## Supplementary Material References

1. H. Li. Seqtk: a fast and lightweight tool for processing FASTA or FASTQ sequences, 2013. Found at: https://github.com/lh3/seqtk.git

2. Edgar RC. MUSCLE: multiple sequence alignment with high accuracy and high throughput. Nucleic Acids Res. 2004 Mar 19;32(5):1792–7. doi: 10.1093/nar/gkh340. PMID: 15034147; PMCID: PMC390337.

3. Tamura K., Stecher G., and Kumar S. (2021). MEGA 11: Molecular Evolutionary Genetics Analysis Version 11. Molecular Biology and Evolution 10.1093/molbev/msab120.

4. Emmanuel Paradis, Klaus Schliep, ape 5.0: an environment for modern phylogenetics and evolutionary analyses in R, *Bioinformatics*, Volume 35, Issue 3, February 2019, Pages 526–528, 10.1093/bioinformatics/bty633

5. Galili T (2015). “dendextend: an R package for visualizing, adjusting, and comparing trees of hierarchical clustering.” Bioinformatics. doi:10.1093/bioinformatics/btv428.

6. Wilkinson SP, Davy SK (2018). *phylogram: an R package for phylogenetic analysis with nested lists*, volume 3. doi:10.21105/joss.00790, http://joss.theoj.org/papers/10.21105/joss.00790.

7. Larkin MA, Blackshields G, Brown NP, Chenna R, McGettigan PA, McWilliam H, Valentin F, Wallace IM, Wilm A, Lopez R, Thompson JD, Gibson TJ, Higgins DG. (2007). Clustal W and Clustal X version 2.0. Bioinformatics, 23, 2947–2948.

8. Nei M. and Kumar S. (2000). Molecular Evolution and Phylogenetics. Oxford University Press, New York.

9. Jain, C., Rodriguez-R, L.M., Phillippy, A.M., et al. High throughput ANI analysis of 90K prokaryotic genomes reveals clear species boundaries. Nat Commun 9, 5114 (2018). 10.1038/s41467-018-07641-9

10. Ivica Letunic, Peer Bork, Interactive Tree Of Life (iTOL) v5: an online tool for phylogenetic tree display and annotation, *Nucleic Acids Research*, Volume 49, Issue W1, 2 July 2021, Pages W293– W296, 10.1093/nar/gkab301

